# MYPT1 O-GlcNAcylation Dictates Timely Disjunction of Centrosomes

**DOI:** 10.1101/795674

**Authors:** Caifei Liu, Yingxin Shi, Xuewen Liu, Zhikai Xiahou, Jie Li, Zhongping Tan, Xing Chen, Jing Li

## Abstract

The role of O-linked N-acetylglucosamine (O-GlcNAc) modification in the cell cycle has been enigmatic. Previously, both O-GlcNAc transferase (OGT) and O-GlcNAcase (OGA) disruption have been shown to derail the mitotic centrosome numbers, suggesting that mitotic O-GlcNAc oscillation needs to be in concert with mitotic progression to account for centrosome integrity. Here we attempted to address the underlying molecular mechanism by both chemical approaches and biological assays, and observed that Thiamet-G (OGA inhibitor) incubation strikingly elevated centrosomal distances, suggestive of premature centrosome disjuction. These aberrancies could be overcome by inhibiting Polo-like kinase 1 (Plk1), a mitotic master kinase. Plk1 inactivation is modulated by the Myosin Phosphatase Targeting Subunit 1 (MYPT1)-Protein Phosphatase 1 cβ (PP1cβ) complex. Interestingly, MYPT1 is abundantly O-GlcNAcylated and the modified residues have been detected in a recent O-GlcNAc profiling screen utilizing chemoenzymatic labeling and bioorthogonal conjugation. We demonstrate that MYPT1 is O-GlcNAcylated at T577, S585, S589 and S601, which antagonizes CDK1-dependent phosphorylation at S473, subsequently attenuating the association between MYPT1 and Plk1, and promoting PLK1 activity. Thus under high O-GlcNAc, Plk1 is untimely activated, conducive to inopportune centrosome separation and disrupting the cell cycle. We propose that too much O-GlcNAc is equally deleterious as too little O-GlcNAc, and a fine balance between the OGT/OGA duo is indispensible for successful mitotic divisions.

## INTRODUCTION

The centrosomes are the primary microtubule-organizing centers that nucleate the mitotic spindle apparatus to ensure subsequent faithful sister chromatid segregation during mitosis. The centrosome cycle is tightly coordinated with other cell cycle events ^1^, and its aberrancy could culminate in chromosome segregation defects and aneuploidy ^2^. The entire centrosome cycle encompasses centrosome duplication during S phase, disjunction in late G2 and further separation during prophase or prometaphase, and eventual segregation into the two daughter cells.

Centrosomes duplicate in concomitant with DNA replication, after which the sister centrosomes are glued together by two proteinaceous linkers, c-Nap1 and rootletin ^3,4^, as well as other components such as Cep68, Cep215 and LRRC45 ^5^. C-Nap1, a large coiled-coiled protein, links rootletin to the centrioles so that the centrosome pair is joined by fibrous polymers ^3^. In late G2, the Never In Mitosis (NIMA)-related serine/threonine kinase Nek2A phosphorylates and displaces c-Nap1 and rootletin, leading to disjoint centrosomes ^6^. Centrosomal accumulation of Nek2A is mediated by the Hippo pathway, among which sterile 20-like kinase 2 (Mst2) and Salvador (Sav1) play critical roles. In particular, Mst2 phosphorylates and activates Nek2A^7^. Upstream of Mst2 is the mitotic master kinase, Polo-like kinase 1 (PLK1) ^8^.

Following centrosome disjunction, the kinesin Eg5 accounts for centrosome positioning in the beginning of M phase. Cyclin-dependent kinase 1 (Cdk1) phosphorylates and activates Eg5 at T927 by stimulating the engagement between Eg5 and microtubules ^9^. Independently, centrosomal localization of Eg5 requires PLK1 ^10^, which activates the NIMA-family kinase Nek9, leading to Eg5 phosphorylation at S1033 by the Nek9/6/7 complex ^11^. Phosphorylated Eg5 then binds centrosomal Targeting Protein for Xenopus kinesin-like protein 2 (TPX2), which is also mediated by Nek9 ^12^. Besides mitosis, Eg5 also governs centrosome dynamics during interphase ^10^. Hence, the centrosomal role of Plk1 is two-fold: centrosome disjunction via the PLK1-Mst2-Nek2A signaling cascade, and centrosome separation through PLK1-Nek9/6/7-Eg5^13^.

Besides centrosomes, PLK1 also orchestrates a multitude of cell cycle events, including replication, mitotic entry, chromosome segregation and cytokinesis ^4,14–16^. It contains an N-terminal kinase domain and a C-terminal polo-box binding domain (PBD). Phosphorylation of PLK1 at T210 at the T-loop is mediated by the Aurora A-Bora complex ^17^, resulting in dissociation of PBD from the kinase domain and thus activating PLK1. Dephosphorylation of PLK1 is modulated by the protein phosphatase 1 cβ (PP1cβ), targeted by the myosin phosphatase targeting subunit 1 (MYPT1) ^18^. Specifically, CDK1 phosphorylates MYPT1 at Ser473, creating a binding pocket between MYPT1 and the PBD of PLK1. Subsequently MYPT1 recruits PP1cβ to dephosphorylate pT210 of PLK1^18^. Such interaction at the kinetochore destabilizes kinetochore-microtububle attachments ^19^. Besides phosphorylation, PLK1 is also methylated at K209 ^20,21^, which vies with pT210 and hence blocking Plk1 activity.

Due to the vital role of PLK1 in mitosis, MYPT1 is subject to multifaceted regulations: the Hippo pathway kinase LATS1/WARTS phosphorylates MYPT1 at S445 to inactivate PLK1 ^22^; optineurin, another phosphatase, promotes MYPT1 activity ^23^; checkpoint kinase 1 (CHK1) phosphorylates MYPT1 at S20, and enhances MYPT1-PP1cβ binding ^24^; checkpoint kinase 2 (CHK2) phosphorylates MYPT1 at S507 to attenuate pS473 ^25^.

Previous investigations have identified that MYPT1 is also subject to O-linked N-acetylglucosamine (O-GlcNAc) modifications ^26^. O-GlcNAcylation is an emerging post-translational modification (PTM) that integrates the metabolic signals with transcription, nutrient sensing, stress responses and cell cycle events ^27,28^. It is catalyzed by the sole transferase O-GlcNAc transferase (OGT), and reversed by the only O-GlcNAcase (OGA) ^27^. Chemical inhibitors of OGT [acetyl-5S-GlcNAc (5S-G)] and OGA [Thiamet-G (TMG)] have been developed to interogate various biological processes^29^. During the cell cycle, O-GlcNAcylation levels fluctuate as the cells proceed through different stages ^30^. In particular, both OGT and OGA overproduction results in multipolar spindles ^31^. However, myriad targets of O-GlcNAc and its quintessential functions remain largely unexplored. Here we identify the O-GlcNAc modified residues of MYPT1. We show that O-GlcNAcylation of MYPT1 antagonizes pS473, and results in its dissociation from PLK1. Elevated O-GlcNAc levels thus fuel PLK1 activity towards centrosomes and render ill-timed centrosome separation, disrupting the mitotic cell cycle.

## RESULTS

### O-GlcNAc promotes aberrant centrosome separation via PLK1

Previously, both OGT and OGA overproduction has been linked with multi-polar spindle ^31^. We sought to identify whether O-GlcNAc could also be linked with centrosome dynamics. Strikingly, when HeLa cells were treated with TMG (OGAi), the inter-centrosomal distance was significantly augmented four-fold (Fig. 1A), reminiscent of the phenotype of Nek2A overexpression or over-activation ^4,32^. As the centrosome cycle is tightly governed by PLK1, we attempted to inhibit PLK1. When BI2536 (PLK1i) was adopted in conjuncture with TMG, the centrosomal distances shortened considerably (Fig. 1A-C). When BI2536 was utilized alone (Fig. 1A), the cells reduced centrosomal distances as previously reported ^4^. These cytological studies suggest that high O-GlcNAc culminates in premature centrosomal separation, probably via PLK1.

**Figure 1.**
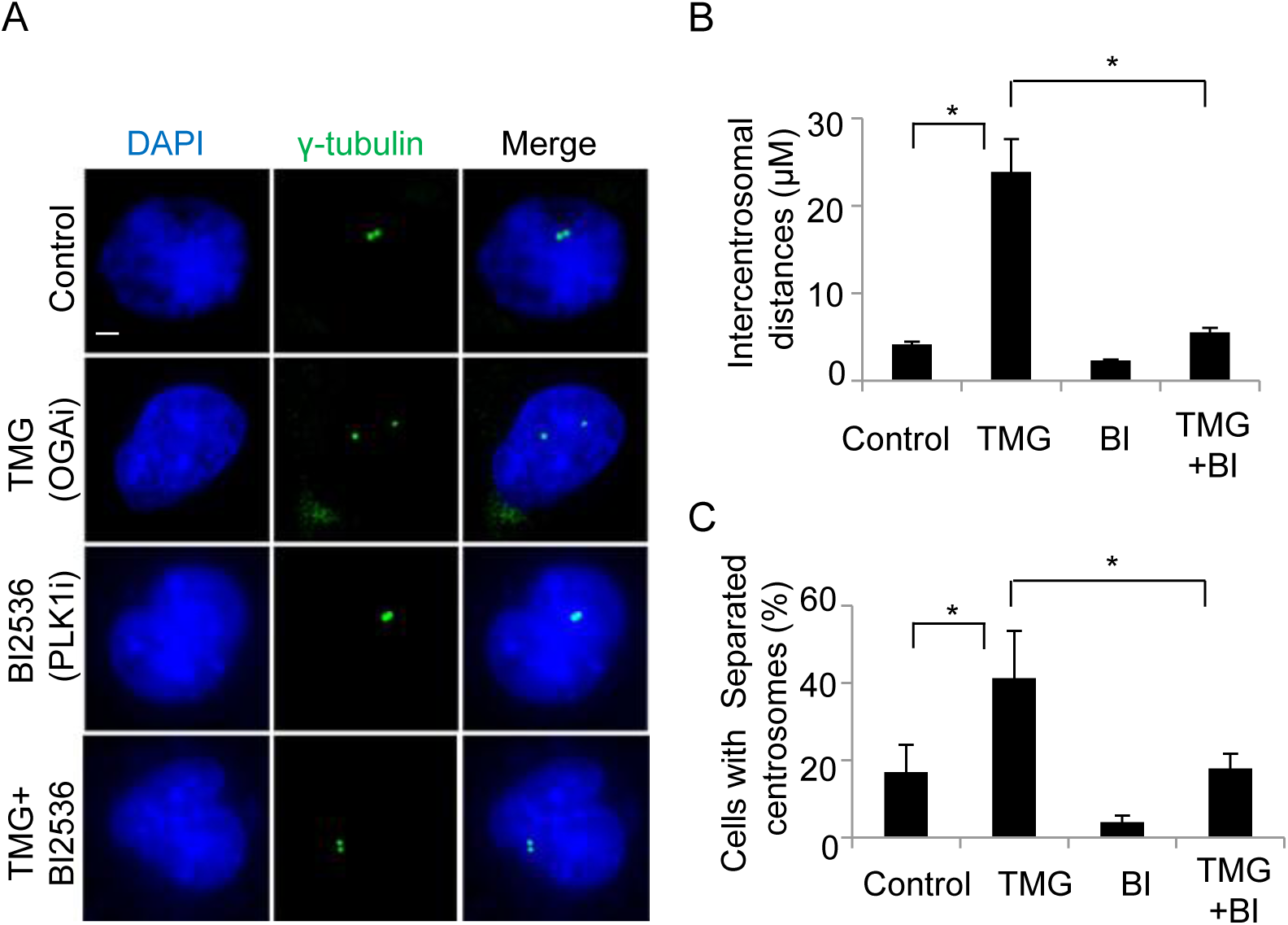
Elevated O-GlcNAc levels leads to aberrant centrosome separation via PLK1. (A) HeLa cells were treated with TMG, BI2536 or TMG + BI2536, then stained with anti-γ-tubulin antibodies and DAPI. Scale bar, 10 μM. (B) Quantitation of inter-centrosomal distances in (A). More than 25 cells were counted for each experiment. The data represent mean ± S. D. of three independent experiments. Asterisks indicate significant difference as determined by t-test (p1-2=0.005, p2-4=0.008). (C) Quantitation of percent of cells with separated centrosomes in (A). Asterisks indicate significant difference as determined by t-test (p1-2=0.02, p2-4=0.02)

### MYPT1 is O-GlcNAcylated at T577, S585, S589 and S601

Previous investigation has identified the inactivating phosphatase of PLK1 is PP1cβ, which is targeted by MYPT1 ^18^. Intriguingly, MYPT1 is O-GlcNAcylated ^26^. Therefore we reasoned that O-GlcNAc might exert its effect through MYPT1.

First, we validated the interaction between MYPT1 and OGT through biochemical assays. As shown in Fig. 2A, GST-OGT pulled-down HA-MYPT1 from cell extracts. Then both OGT and MYPT1 proteins were purified from *E. coli.* Upon incubation, His-OGT pulled-down GST-MYPT1 (Fig. 2B), suggesting that the interact is direct.

**Figure 2.**
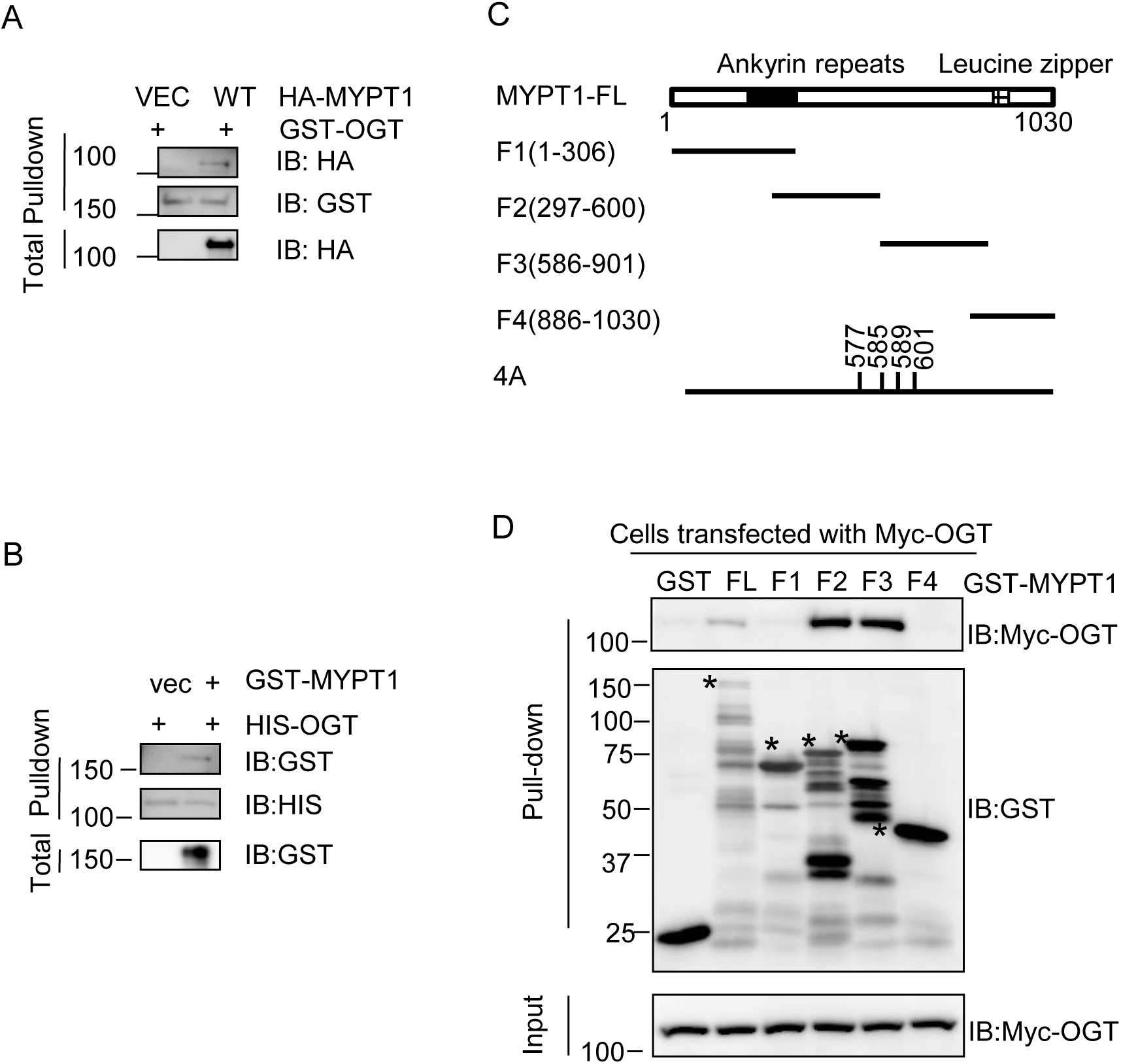
OGT interacts with the central region of MYPT1. (A) Recombinant GST-OGT proteins were incubated with HA-MYPT1-transfected cell lysates. (B) His-OGT and GST-MYPT1 proteins were incubated together and then subject to pulldown assays as indicated. (C) A diagram showing MYPT1 constructs used in this study. Full-length (FL), F1(1-306), F2(297-600), F3(586-901) and F4(886-1030) were previously described ^24^. 4A denotes T577AS585AS589AS601A. (D) Recombinant GST-MYPT1-FL, F1, F2, F3 and F4 proteins were purified from bacteria, and incubated with extracts from 293T cells transfected with Myc-OGT. Asterisks demarcate corresponding proteins.

Then we mapped which domain of MYPT1 interacts with OGT. As MYPT1 is a relative large protein, we constructed several fragments of MYPT1 as previously described: F1 (1-306), F2 (297-600), F3 (586-901) and F4 (886-1030) (Fig. 2C) ^24^. To investigate which fragment interacts with OGT, recombinant full-length (FL) and F1-F4 MYPT1 proteins were utilized in pulldown experiments, and the FL, F2 and F3 MYPT1 pulled-down Myc-OGT (Fig. 2D), suggesting that the potential modification sites could be residing in F2 and F3.

A recent quantitative proteomic analysis of protein O-GlcNAc sites using an isotope-tagged cleavable linker (isoTCL) strategy identified the potential O-GlcNAc sites of MYPT1 to be T577, S585, S589 and S601 ^33^ (Fig. 3 A-D), all of which locate on F2 and F3. We constructed the T577A/S585A/S589A/S601A (4A) mutant accordingly and assessed its effect. When HA-MYPT1-wild type (WT) and 4A plasmids were transfected into cells, the 4A mutant significantly abrogated O-GlcNAcylation (Fig. 4A), suggesting that these four amino acids are major O-GlcNAc sites. Considering that MYPT1 is abundantly O-GlcNAcylated, and other proteomic screens have also identified extra glycosylation sites ^34^, our results do not exclude the possibility that there could be more O-GlcNAcylated residues on MYPT1.

**Figure 3.**
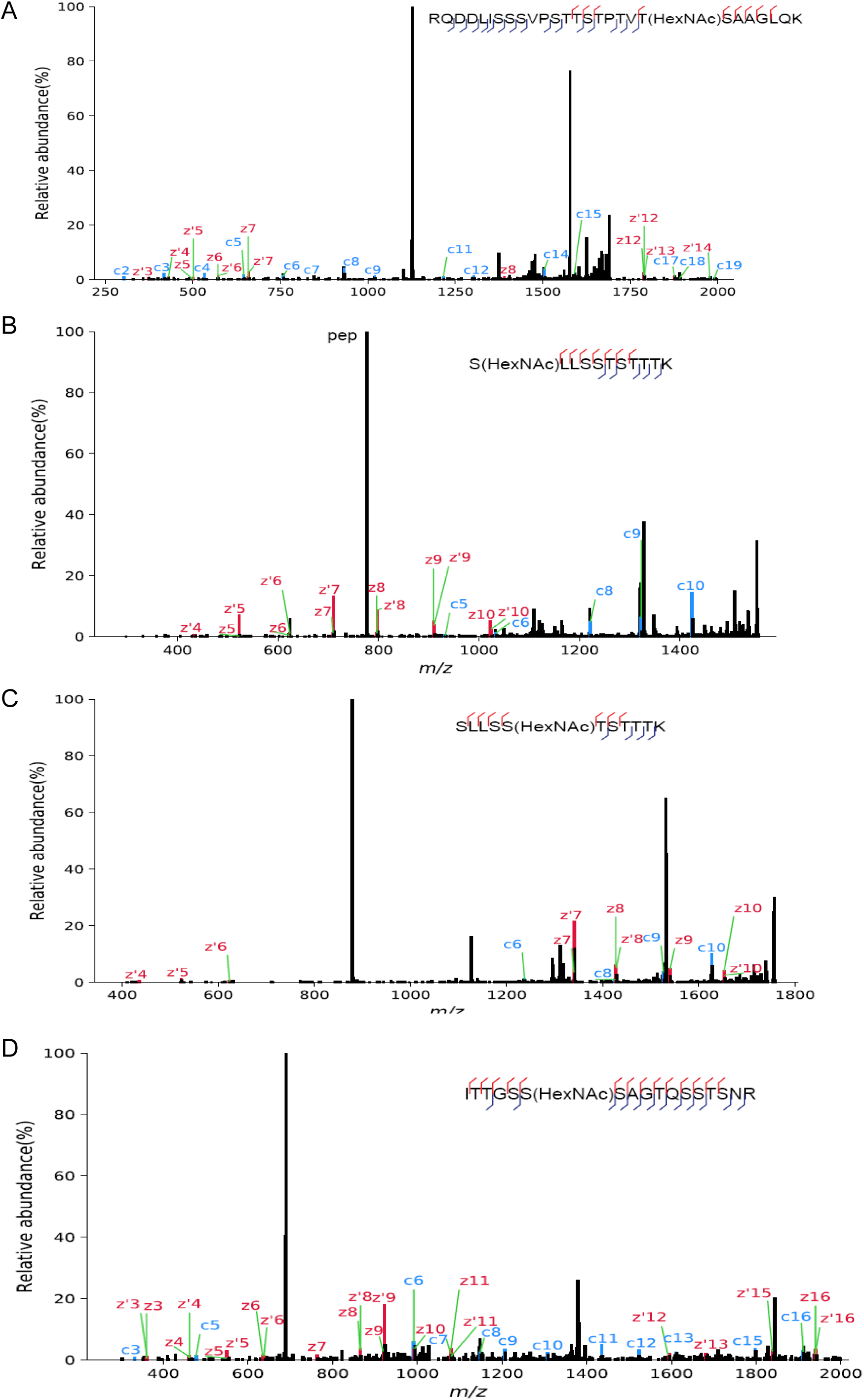
MYPT1 is O-GlcNAcylated at T577, S585, S589 and S601. (A-D) Electron Transfer Dissociation (ETD) mass spectrometry combined with chemo-enzymatic labeling identified that T577S585 S589S601 are O-GlcNAcylated ^33^.

**Figure 4.**
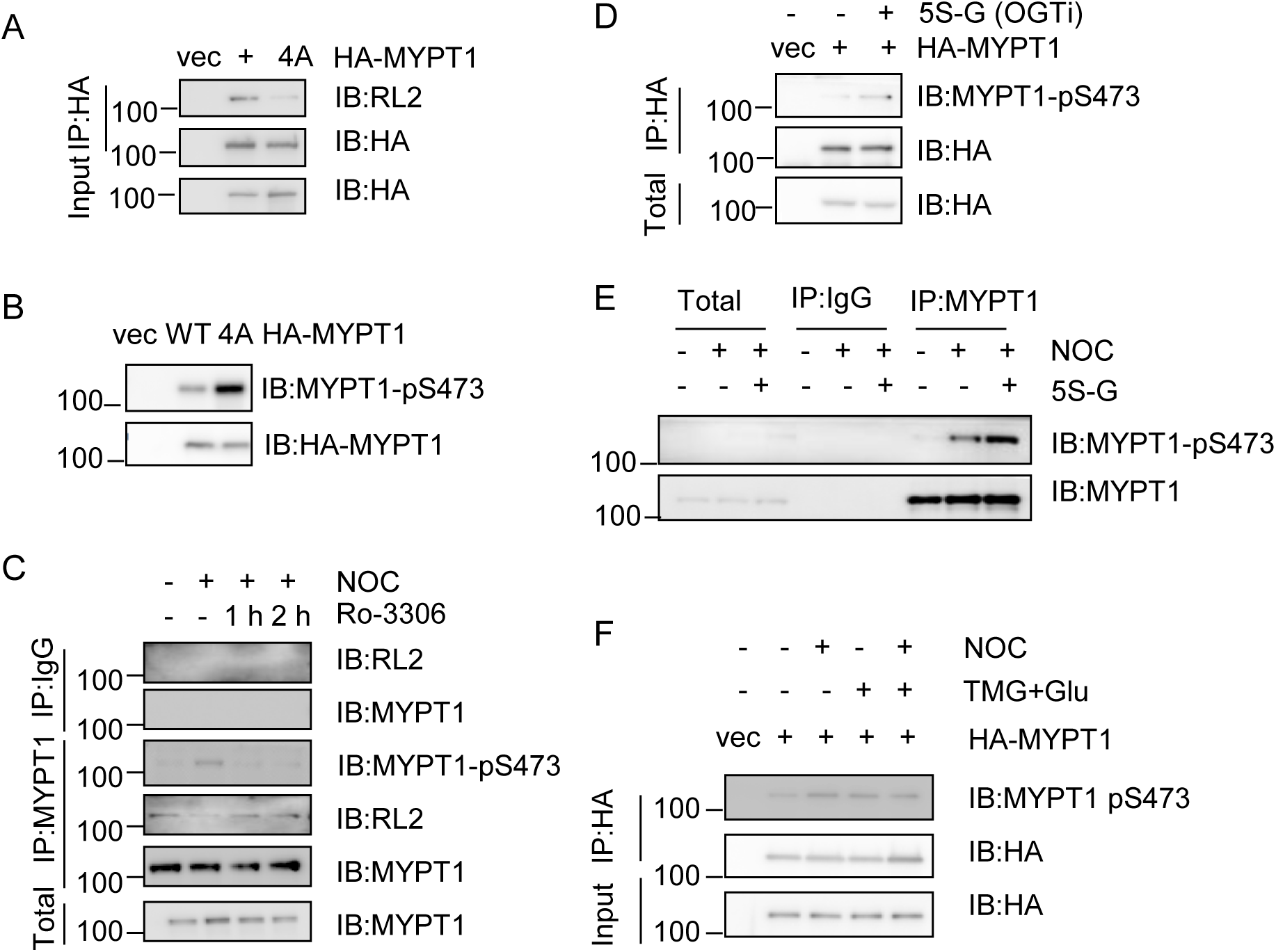
O-GlcNAcylation of MYPT1 antagonizes CDK1-dependent phosphorylation at S473. (A) MYPT1-WT and 4A plasmids together with Myc-OGT or empty vectors were transfected into 293T cells and then blotted with antibodies indicated. (B) Cells were transfected with HA-MYPT1-WT or 4A plasmids, and then the lysates were IBed with antibodies indicated. (C) Cells were treated with Noc, or Noc with Ro-3306 for the time indicated. (D) HeLa cells were transfected with HA-MYPT1, treated or untreated with 5S-G (OGT inhibitor). (E) Cells were treated with Noc, or Noc + 5S-G. (F) Cells were transfected with MYPT1-WT plasmids, and then treated with Noc, or Noc plus TMG + Glu as indicated.

### O-GlcNAcylation of MYPT1 antagonizes CDK1-dependent phosphorylation at S473

Since CDK1 phosphorylates MYPT1 at S473 during mitosis and creates a binding motif between MYPT1 and the PBD of PLK1^18^, we surmised that O-GlcNAc of MYPT1 might interplay with pS473. To address this possibility, we used a phospho-specific antibody targeting pS473 that has been previously described and utilized ^25^.

Then the WT and 4A plasmids are compared for the pS473 levels, and it is significantly bolstered in the 4A mutant (Fig. 4B). When Noc was used to synchronize cells in the M phase, O-GlcNAc levels decreased while pS473 levels arose (Fig. 4C). As pS473 is mediated by CDK1, we adopted RO-3306 again, and observed that RL2 levels decreased while pS473 levels increased in the RO-3306-treated cells (Fig.4C). This is consistent with our conjecture that O-GlcNAc antagonizes pS473. Lastly, we utilized the 5S-G inhibitor for OGT ^29^, and 5S-G treatment substantially boosted pS473 levels of transfected HA-MYPT1(Fig. 4D). We also examined the effects of 5S-G on endogenous MYPT1. Noc treatment enhanced pS473 levels, and Noc plus 5S-G elevated pS473 markedly (Fig. 4E). In contrast, glucose plus TMG (TMG+Glu) treatment during Noc would down-regulate pS473 compared to Noc alone (Fig. 4F). Taken together, O-GlcNAc of MYPT1 attenuates pS473.

### O-GlcNAcylation inhibits MYPT1-PLK1 association

Since pS473 promotes MYPT1-PLK1 association ^18^, we then explored the effect of O-GlcNAc on the interaction between MYPT1 and PLK1 by treating the cells with TMG+Glu to enhance O-GlcNAc ^35,36^. As shown in Fig. 5A, Noc increased PLK1-MYPT1 association discernably as reported ^18^, but TMG+Glu together with Noc obliterated PLK1-MYPT1 affinity.

**Figure 5.**
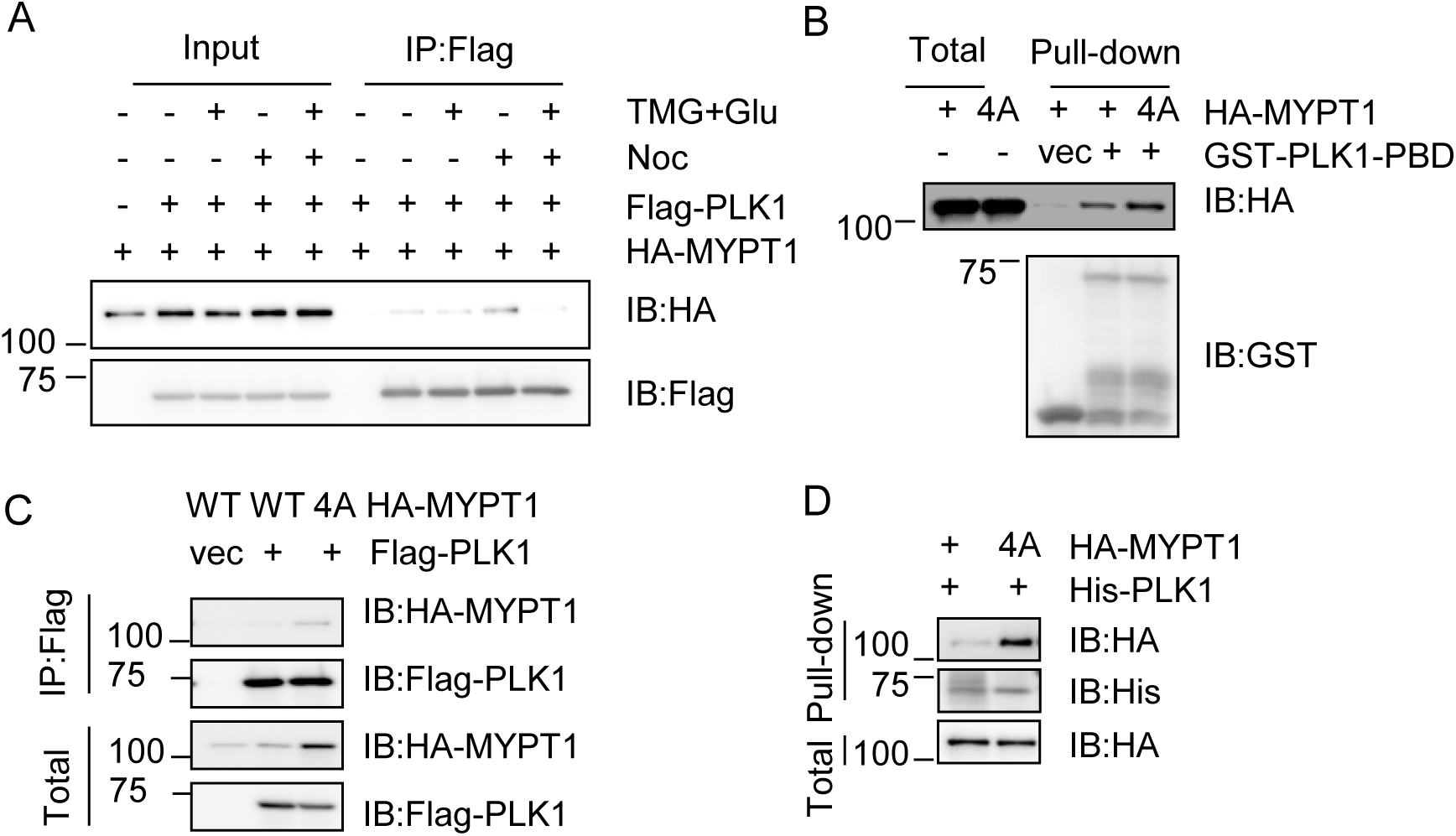
O-GlcNAcylation of MYPT1 attenuates the interaction between MYPT1 and PLK1. (A) 293T cells were transfected with Flag-PLK1 and HA-MYPT1, treated or not treated with Noc, TMG + Glu, respectively, then subject to IP and IB as indicated. (B) GST-PLK1-PBD proteins were purified from bacteria. Cells were transfected with HA-MYPT1-WT or 4A, then the cell lysates were subject to GST-PLK1-PBD pulldown assays. (C) Cells were transfected with FLAG-PLK1 together with HA-MYPT1-WT or 4A, then subject to IP and IB as indicated. (D) Cells were transfected with HA-MYPT1-WT or 4A, and then cell extracts were utilized in His-PLK1 pulldown assays.

As phosphorylated MYPT1 binds with PLK1-PBD ^18^, we adopted GST pulldown experiments using PLK1-PBD, and GST-PLK1-PBD modestly increased binding with HA-MYPT1-4A (Fig. 5B). Then we employed FL-PLK1. When we directly utilized the 4A mutant to coIP PLK1, the interaction between MYPT1 and PLK1 substantially upregulated (Fig. 5C).When His-PLK1 was applied in pulldown assays, 4A again manifested more robust association with PLK1(Fig. 5D). In sum, the binding between PLK1 and MYPT1 was abolished during high O-GlcNAc.

### O-GlcNAcylation of MYPT1 enhances PLK1 activity

As the MYPT1 associates PLK1 to target PP1cβ to dephosphorylate and deactivate PLK1 ^18^, stronger affinity could signify less activity. We took advantage of the IP-phosphatase assay to examine PLK1 activity ^22,24^. Cells were transfected with Flag-MYPT1 and treated with Noc. Cells were also supplemented with TMG + Glu to enrich for O-GlcNAc or not treated. When the anti-FLAG immunoprecipitates were incubated with recombinant PLK1, the relative low O-GlcNAc group efficiently dephosphorylated PLK1, as detected by IB with PLK1-pT210 antibodies, but not the high O-GlcNAc group (Fig. 6A).

**Figure 6.**
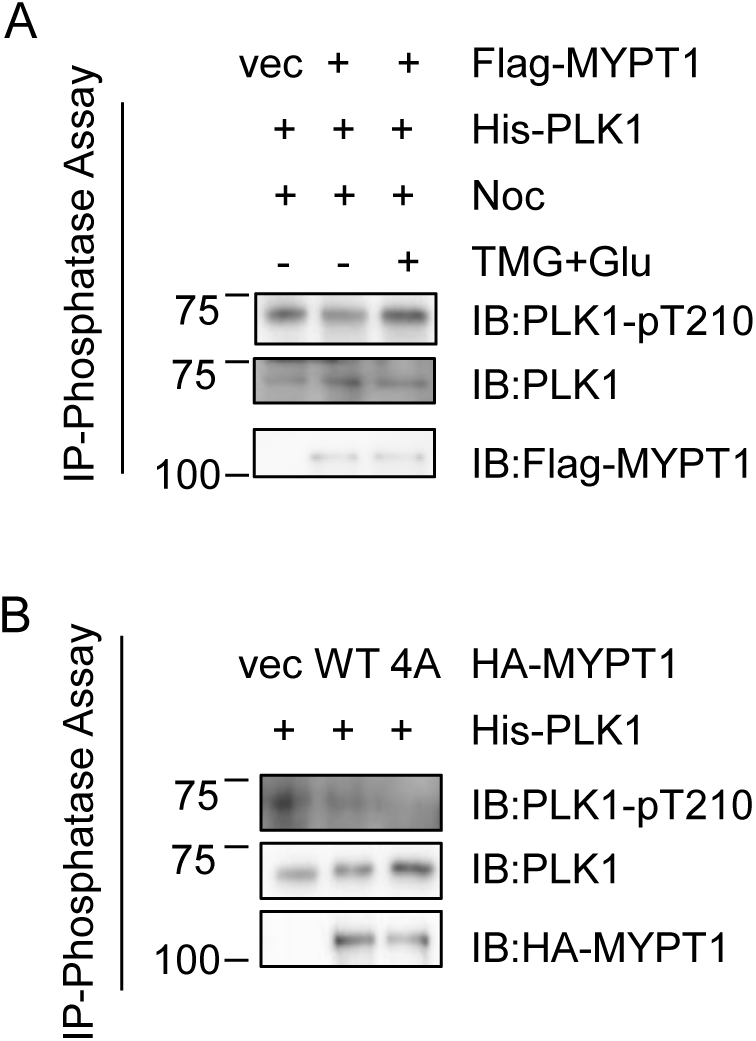
O-GlcNAcylation of MYPT1 promotes PLK1 activity. (A) IP-phosphatase assays. U2OS cells were transfected with Flag-MYPT1, synchronized to mitosis with Noc, then treated with TMG + Glu or left untreated. The anti-Flag immunoprecipitates were then incubated with recombinant His-PLK1. (B) IP-phosphatase assays using the MYPT1-WT and −4A mutants without Noc treatment.

MYPT1-4A mutants were then directly exploited in the IP-phosphate assay. In the absence of Noc, MYPT1-WT decreased PLK1 pT210 levels, and the MYPT1-4A completely abolished PLK1-pT210 levels (Fig. 6B). This is consistent with our results in Fig. 5C-D that MYPT1-4A partners with PLK1 in the absence of Noc treatment. Collectively, our biochemical assays suggest that O-GlcNAcylated MYPT1 disjoins PLK1 and promotes its kinase activity.

### MYPT1-4A Suppresses the TMG-induced centrosome disjunction defects

Since the aforementioned results suggest that MYPT1 O-GlcNAcylation is a pivotal regulator in centrosome separation, we undertook sh*MYPT1* to knockdown endogenous MYPT1 (Fig. 7A), so that the effects of MYPT1-4A could be directly measured and observed after TMG incubation. As shown in Fig. 7B, the premature centrosome separation phenotype is discernable in the sh*MYPT1* cells that bears MYPT1-WT plasmids. But in the cells transfected with MYPT1-4A plasmids, the aberrancy is suppressed (Fig. 7C), in line with previous reports that PLK1 sequestration culminates in duplicated but unseparated centrosomes^37,38^. Taken together, the 4A mutant fails to show the untimely centrosome separation phenotype, probably due to PLK1 suppression.

**Figure 7.**
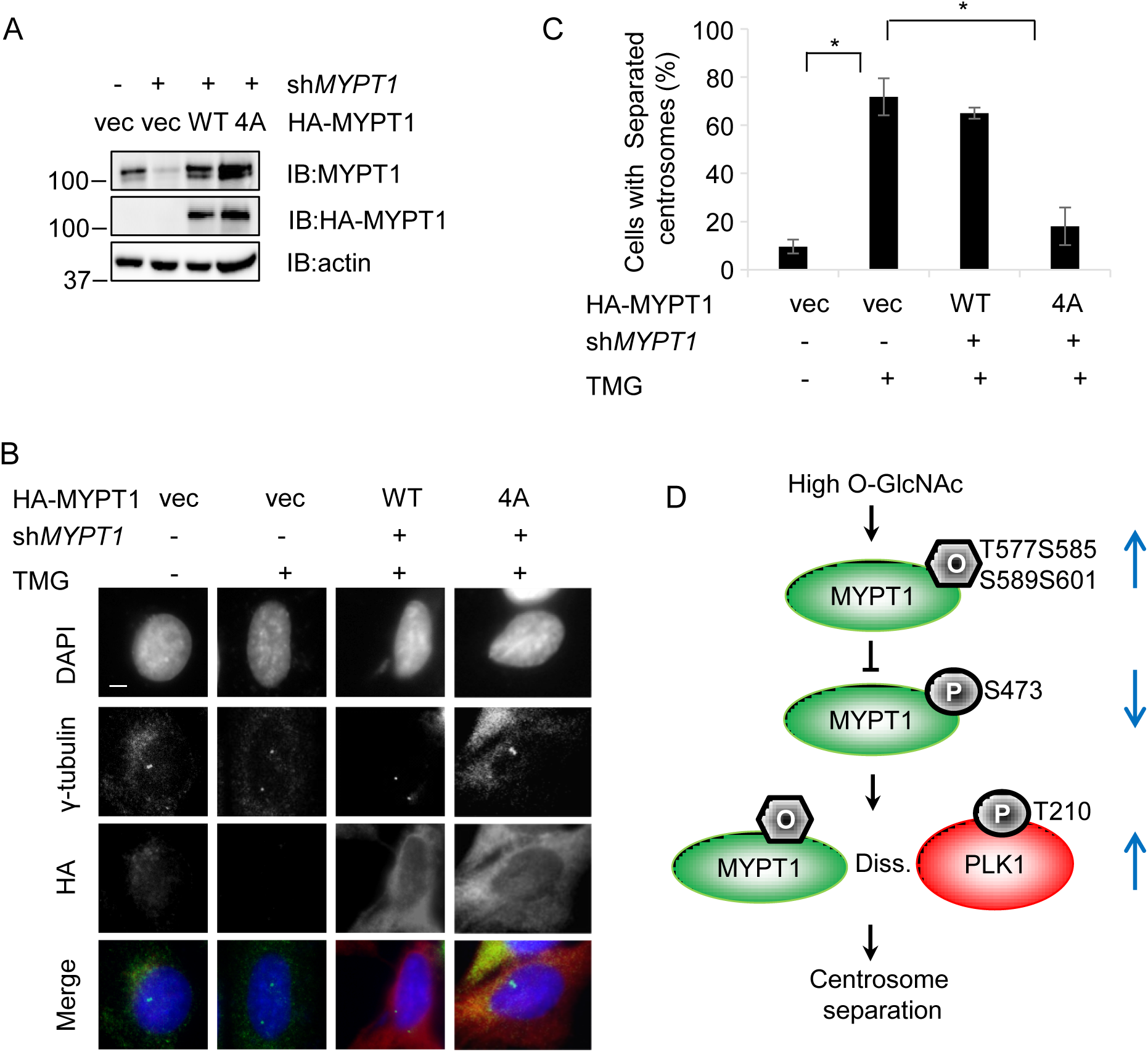
MYPT1 overproduction overrides the centrosome disjunction defects induced by TMG. (A) Lenti viruses encoding vectors or sh*MYPT1* was introduced into HeLa cells, together with HA-MYPT1-WT or −4A plasmids. The cellular lysates were IBed with the antibodies indicated. (B) Cells in (A) were subject to indirect IF using the antibodies indicated. (C) Quantitation of percent of cells with separated centrosomes in (B). Asterisks indicate significant difference as determined by t-test (p1-2=0.0002, p2-3=0.22, p2-4=0.001). (D) We propose that MYPT1 is O-GlcNAcylated at T577 S585 S589 S601, which antagonizes CDK1-dependent phosphorylation at S473 and MYPT1-PLK1 interaction. By disjoining PLK1 from the MYPT1/PP1cβ complex, PLK1 activity is elevated, thus promoting centrosome separation.

## DISCUSSION

In this study, we identify that O-GlcNAc regulates centrosome separation through the MYPT1-PLK1 complex. We pinpoint the major O-GlcNAcylated sites of MYPT1 to be S564, S566, T570 and S578 in human cells, and further delineate that O-GlcNAc antagonizes pS473, hinders MYPT1-PLK1 association, thus boosting PLK1 activity and hence centrosome disjunction (Fig. 7D). When MYPT1 fails to be O-GlcNAcylated, as in the MYPT1-4A mutant, PLK1 activity is dampened (Fig. 6B).

MYPT1 is one of the most abundant O-GlcNAcylated proteins, and its modification sites have been unveiled time and again in distinct proteomic studies ^34,39,40^. Perhaps O-GlcNAc sites might not be conserved between humans and mice. MYPT1 is found to be O-GlcNAcylated at S564, S566, T570 and S578 by mass spectrometry in the mouse brain ^40^, but our data show in HeLa cells O-GlcNAc occurs in T577, S585, S589 and S601. Previously, p53 is identified to be O-GlcNAcylated at S149 ^41^, which is not conserved in mice either. The same also holds true for phosphorylation. For instance, Ataxia-telangiectasia mutated (ATM), a vital sensor protein for DNA damage signaling, is phosphorylated at S1981 and then activated in humans, but mutation of S1987 (the mouse equivalent) does not hinder ATM function in mice^42,43^. Thus extrapolating data across species needs extra caution, as the function and sites of PTMs could be context-dependent.

Along the same vein, there could be more O-GlcNAc sites on MYPT1, as the proteomic studies were carried out under disparate circumstances and using different click chemistry methodologies ^33,34,39,40^. As the O-GlcNAc modification is highly dynamic, distinct sites could be modified in response to environmental cues.

O-GlcNAc could have multi-faceted effects on the centrosome. Both OGT and OGA overexpression result in multi-polar spindles, which could be repressed by TMG treatment ^31^. NuMA (nuclear mitotic apparatus protein) is indispensible for spindle pole formation and regulates spindle pole cohesion ^44,45^. NuMA is O-GlcNAcylated and its localization was led astray by OGT overexpression^46^. Further investigations reveal that O-GlcNAcylated NuMA interacts with Galectin-3, which is a prerequisite for mitotic spindle cohesion and proper NuMA localization ^47^. Here we reveal that centrosome dynamics is also governed by O-GlcNAcylation levels. As the centrosome is pivotal for the mitotic process, O-GlcNAc is bound to modulate other aspects of centrosome function.

Our results indicate that O-GlcNAcylated MYPT1 attenuates interaction with PLK1 and thus promotes PLK1 activity. It is intriguing that overall PLK1 pT210 levels remain unaltered in OGT or OGA overproduction cells ^31,46^. We did not detect discernable difference either, in cells supplemented with TMG plus Glucose (data not shown). This may seem paradoxical at first, but considering the versatile roles of PLK1 during mitosis ^4,14–16^, we could entertain the possibility that only a small pool of PLK1 is regulated by MYPT1. First, although the overall activity of PLK1 is upregulated during mitosis, CDK1 actually dampens PLK1 activity via MYPT1 in a mitosis-specific fashion ^18^. Secondly, the pool of PLK1 responsible for kinetochore-microtubule attachment actually contains low PLK1 kinase activity during metaphase so that microtubules could be dynamic ^48^. Therefore, irrespective of the overall elevation of mitotic activity, the mitotic master kinase - PLK1 is perhaps indeed fine-tuned in space and time. And O-GlcNAc, could be the sweet icing on the cake.

## EXPERIMENTAL PROCEDURES

### Cell culture, antibodies and plasmids

Cells were purchased from ATCC. Antibodies were as follows: anti-MYPT1 (Bethyl, # BL3866), anti-PLK1 (Zymed, #37-7100), PLK1-pT210 (BD Pharmingen, #558400). Antibodies against pS473 were prepared as described before ^25^ and manufactured by Beijing B&M Biotech Co., Ltd.. *MYPT1* plasmids were described before ^49^. *MYPT1-4A* plasmids were generated using specific primers (sequences available upon request) following the manufacturer’s instructions (QuickChange II, Stratagene). His-OGT was from Dr. Yue Wang (Peking Univ.). The following shRNA target sequences were used: sh*MYPT1:* GTAACCCAGTGGACCATAATT.

### Immunoprecipitation (IP) and Immunoblotting (IB) assays

IP and IB experiments were performed as described before ^50^. The following primary antibodies were used for IB: anti-β-actin (1:10000), anti-HA (1:1000), and anti-FLAG M2 (Sigma) (1:1000), anti-Myc (1:1000), anti-PLK1 (1:1000), anti-MYPT1 (1:1000), PLK1-pT210 (1:500). The IP-phosphatase assay was performed as before ^22,24^.

Peroxidase-conjugated secondary antibodies were from JacksonImmuno Research. Blotted proteins were visualized using the ECL detection system (Amersham). Signals were detected by a LAS-4000, and quantitatively analyzed by densitometry using the Multi Gauge software (Fujifilm). All western blots were repeated for at least three times.

### Cell Culture Treatment

Chemical utilization: Nocodazole (Noc) at 100 ng/ml for 16 hours; Ro 3306 (CDK1 inhibitor) at 2 μM for the time indicated; BI2536 (PLK1 inhibitor) at 100 nM for two hours; Thiamet-G (TMG) (OGA inhibitor) at 5μM for 24 hrs; acetyl-5S-GlcNAc (5S-G) (OGT inhibitor) was used at 100μM (prepared at 50 mM in DMSO) for 24 hrs [64].

### Indirect Immunofluorescence

Indirect immunofluorescence staining was performed as described before ^50^. Dilutions of primary antibodies were 1:1,000 for mouse anti-γ-tubulin. Cell nuclei were stained with DAPI. Quantitation was performed with the software Image J.

## Abbreviations

PTM: post-translational modification
O-GlcNAc: O-linked N-acetylglucosamine
OGT: O-GlcNAc transferase
TMGP: Thiamet-G
PBD: polo-box binding domain
MYPT1: myosin phosphatase targeting subunit 1
Cdk1: Cyclin-dependent kinase 1
PLK1: Polo-like kinase 1
5S-G: acetyl-5S-GlcNAc
PP1cβ: protein phosphatase 1 c β
FL: full-length
ETD: Electron Transfer Dissociation

## Acknowledgements

We thank Dr. Hai-Ning Du (Wuhan Univ.) and the Li laboratory for helpful discussion. This work is supported by the National Natural Science Foundation of China (NSFC) fund (31872720) and Capacity Building for Sci-Tech Innovation - Fundamental Scientific Research Funds (19530050137) to J. L.; NSFC (NOs. 21425204, 21672013 and 21521003) and the National Key Research and Development Projects (NOs. 2016YFA0501500 and 2018YFA0507600) to X.C.; NSFC (91853120) and the National Major Scientific and Technological Special Project of China for "Significant New Drugs Development" (2018ZX09711001-013) to Z. T.

## Competing interests

The authors declare that they have no conflicts of interest with the contents of this article.

## Author contributions

J. L., X.C. and Z.T. conceived the project and analyzed the data. C. L., Y. S., X. L., Z. X. and Jie L. performed all the experiments. All authors reviewed and approved the manuscript.

